# Solubilization of Membrane Proteins using designed protein WRAPS

**DOI:** 10.1101/2025.02.04.636539

**Authors:** Ljubica Mihaljević, David E. Kim, Helen E. Eisenach, Pooja D. Bandawane, Andrew J. Borst, Alexis Courbet, Everton Bettin, Qiushi Liu, Connor Weidle, Sagardip Majumder, Xinting Li, Mila Lamb, Analisa Nicole Azcárraga Murray, Rashmi Ravichandran, Elizabeth C. Williams, Shuyuan Hu, Lynda Stuart, Linda Grillová, Nicholas R. Thomson, Pengxiang Chang, Melissa J. Caimano, Kelly L. Hawley, Neil P. King, David Baker

**Affiliations:** Department of Biochemistry, Institute for Protein Design, University of Washington, Seattle, WA 98195, USA; Departments of Pediatrics, UConn Health, Farmington, CT 06030-3715, USA; Medicine, UConn Health, Farmington, CT 06030-3715, USA; Molecular Biology and Biophysics, UConn Health, Farmington, CT 06030-3715, USA; Division of Infectious Diseases and Immunology, Connecticut Children’s, Hartford CT 06106, USA; Department of Research, Connecticut Children’s Research Institute, Hartford CT 06106, USA; Howard Hughes Medical Institute, University of Washington, Seattle, WA, USA; Centre de Biologie Structurale, University of Montpellier, CNRS, INSERM. Montpellier, France; Kymab, a Sanofi Company, Babraham Research Campus, Cambridge, UK; Parasites and Microbes Programme, Wellcome Sanger Institute, Hinxton, UK; Faculty of Infectious and Tropical Diseases, London School of Hygiene & Tropical Medicine, London, UK

**Author notes:** These authors contributed equally to this work (Ljubica Mihaljević, David E. Kim).

## Abstract

The development of therapies and vaccines targeting integral membrane proteins has been complicated by their extensive hydrophobic surfaces, which can make production and structural characterization difficult. Here we describe a general deep learning-based design approach for solubilizing native membrane proteins while preserving their sequence, fold, and function using genetically encoded *de novo* protein WRAPs (<u>W</u>ater-soluble <u>R</u>Fdiffused <u>A</u>mphipathic <u>P</u>roteins) that surround the lipid-interacting hydrophobic surfaces, rendering them stable and water-soluble without the need for detergents. We design WRAPs for both beta-barrel outer membrane and helical multi-pass transmembrane proteins, and show that the solubilized proteins retain the binding and enzymatic functions of the native targets with enhanced stability. Syphilis vaccine development has been hindered by difficulties in characterizing and producing the outer membrane protein antigens; we generated soluble versions of four *Treponema pallidum* outer membrane beta barrels which are potential syphilis vaccine antigens. A 4.0 Å cryo-EM map of WRAPed TP0698 is closely consistent with the design model. WRAPs should be broadly useful for facilitating biochemical and structural characterization of integral membrane proteins, enabling therapeutic discovery by screening against purified soluble targets, and generating antigenically intact immunogens for vaccine development.

## Main Text

Membrane proteins constitute nearly a third of the proteome (1), are targeted by over half of clinically approved drugs (1,2), and have considerable potential as vaccine targets (3). However, the challenges associated with working with membrane proteins, particularly their extraction and poor stability (4), remain significant barriers to research and drug discovery efforts. Redesigning membrane proteins to be expressed in a soluble form could bypass the need for conventional extraction from the membrane with detergents (5–8). The majority of existing redesign methods rely on modifying native protein sequences by mutating lipid-facing hydrophobic residues to polar amino acids, either through manual manipulation (5) or computational approaches (6–8). Genetic fusion of the amphipathic apolipoprotein AI (ApoA-I) to membrane proteins conferred water solubility without modification of the native membrane protein sequence by shielding the lipid-facing hydrophobic regions through nonspecific hydrophobic interactions (9); however, the flexibility of ApoA-I and the lack of specificity of the interactions with the shielded target could complicate structure determination (9).

We reasoned that new machine learning-based tools for *de novo* protein design could enable the design of custom solubilizing domains (WRAPs) for each membrane protein target. As in the ApoA-I approach, the resultant designed fusion proteins would have a polar exterior and a nonpolar interior complementary to the lipid-facing hydrophobic surface of the target. We set out to explore the use of RFdiffusion, a generative model for *de novo* protein backbone design (10), to generate both alpha-helical and beta-sheet target-specific solubilizing domains.

## *De novo* design of WRAPs

In initial experiments, we found it difficult to guide free diffusion off of the terminus of a target membrane protein using random lengths of amino acids to fully protect all of the lipid-facing hydrophobic regions. Instead, we turned to a two-step approach in which we first generated idealized *de novo* helical and beta-barrel backbones for WRAPs that roughly matched the diameter and height of the membrane-spanning portion of the target protein (Figure 1A). Cylindrical antiparallel helical assemblies with an inner diameter compatible with the membrane-spanning region of the target protein were generated using fold-conditioned RFdiffusion (10) (Figure S1A). To generate beta-barrel backbones, we built ideal backbones parametrically (11), selecting the length and number of beta strands and the barrel shear (the register shift between the first and last strands) to generate backbones with internal diameters and barrel heights to accommodate side chain packing interactions with the target protein while maintaining the interstrand hydrogen bonds of the barrel.

**Fig. 1.**
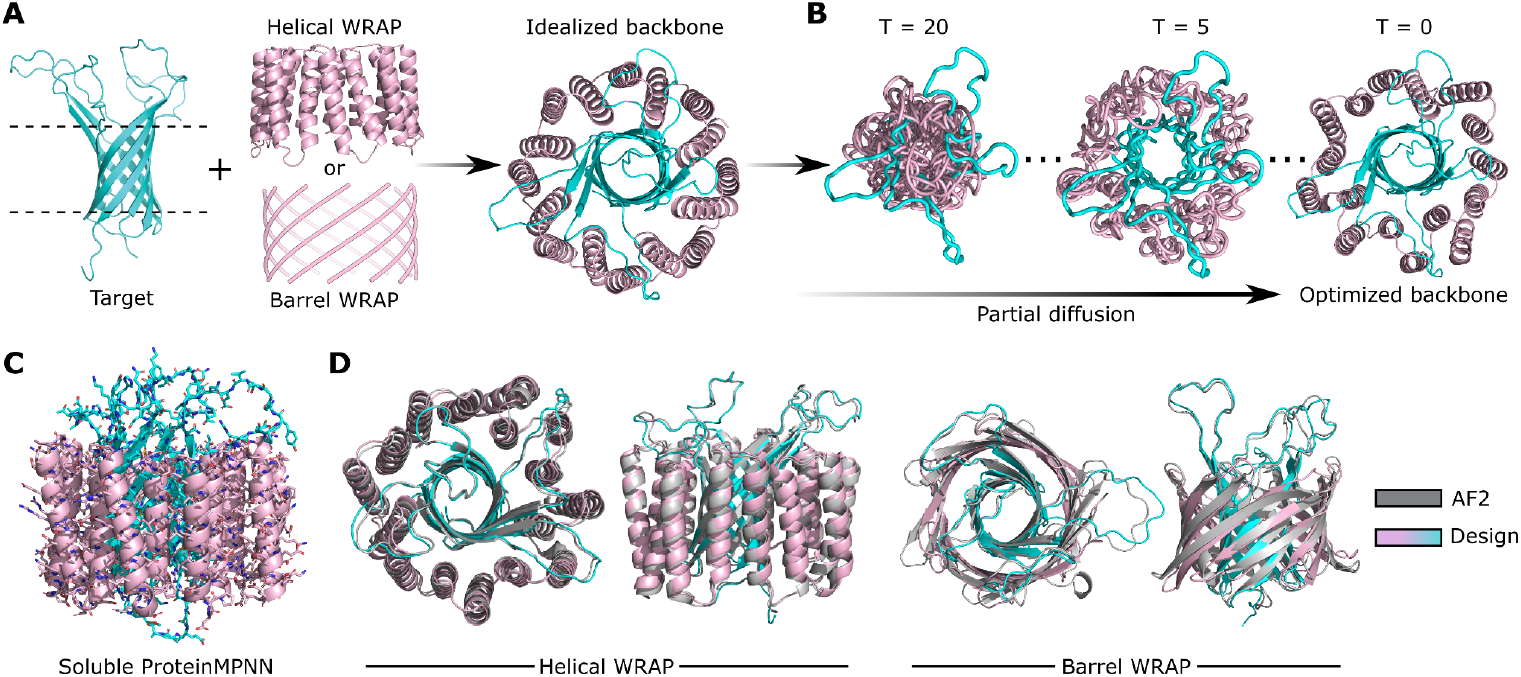
Pipeline for *de novo* design of WRAPs. **(A)** Protein placement and backbone integration: target membrane proteins (e.g., *E. coli* OmpA, cyan) were centrally positioned within WRAP backbones (pink) generated through fold-conditioned RFdiffusion (for helical structures) or parametric design (for barrel structures). An idealized backbone, comprising WRAP as chain A and the target protein as chain B, was established as the starting point for subsequent refinement steps. **(B)** Backbone refinement: Partial diffusion over 20 timesteps (T) was applied to facilitate optimal backbone-to-backbone interactions. **(C)** Sequence design: Sequences were designed for the optimized backbones using soluble ProteinMPNN. **(D)** Structure prediction: AF2-predicted structure of the designs (gray) is overlaid with the WRAP-OmpA design models (pink-cyan). Confidence metrics provided by AF2 guided the selection of designs for experimental validation.

We next refined the designed backbones to be complementary in shape and side chain interactions to the target protein using partial diffusion (12) to closely match the target surface (Figure 1B). The target protein was positioned in the center of the designed helical or barrel WRAP backbones using PyMOL or PyRosetta (13) (Figure 1A). Partial diffusion was then used to mold the *de novo* backbone(s) to the target by partially randomizing the coordinates of the designed backbones and successively removing noise using RFdiffusion conditioned on the target structure. With this conditioning, the resultant optimized backbones are expected to be as complementary in shape to the target as typical protein-protein interfaces in the PDB (Figure 1B). We then designed amino acid sequences for the *de novo* WRAP domains using a variant of ProteinMPNN (Figure 1C). Standard ProteinMPNN (14) was trained from a set of PDB structures that included transmembrane proteins and, as a result, it tends to place nonpolar residues on the exposed surface of long alpha helices. A version of ProteinMPNN specialized for soluble proteins (SolubleMPNN) (15) was therefore used to design sequences for the refined backbones, keeping the amino acid identities but not the side chain conformations of the target membrane protein residues fixed, yielding sequences that direct folding to the desired structure and make specific interactions with the target (Figure 1C). AlphaFold2 (AF2) (16) structure prediction metrics were then used to select promising designs for experimental characterization (Figure 1D; pLDDT (per-residue confidence score) > 85; predicted aligned error of interaction (PAE_i) < 8; root mean square deviation (RMSD) to the design < 1 Å).

## Solubilization of outer membrane beta-barrel proteins (OMPs)

Outer membrane proteins (OMPs) are transmembrane beta-barrel proteins found in diderm bacteria and upon recombinant expression require refolding from inclusion bodies, in addition to purification with detergents (17). As a first test of our method, we sought to solubilize the outer membrane protein A (OmpA) of *E. coli*, a well-characterized 8-stranded transmembrane beta-barrel protein (18). We generated both alpha-helical and beta-barrel WRAPs for OmpA, and expressed the WRAP-OmpA fusion proteins in the *E. coli* cytoplasm. We were able to purify both helical (Figure 2A) and beta-barrel (Figure 2D) WRAP-OmpA proteins from the soluble fraction of *E. coli* lysates. Five out of six N-terminally fused helical WRAP-OmpA designs and one out of six beta barrel WRAP-OmpA C-terminally fused designs were soluble. Size exclusion chromatography (SEC) profiles of representative helical and beta-barrel WRAP-OmpA proteins (Figure 2B and 2E, respectively) were monodisperse with single well-defined peaks, and circular dichroism (CD) spectra and temperature melts (Figure 2C, 2F) showed that both designs were folded and stable at 95°C.

**Fig. 2.**
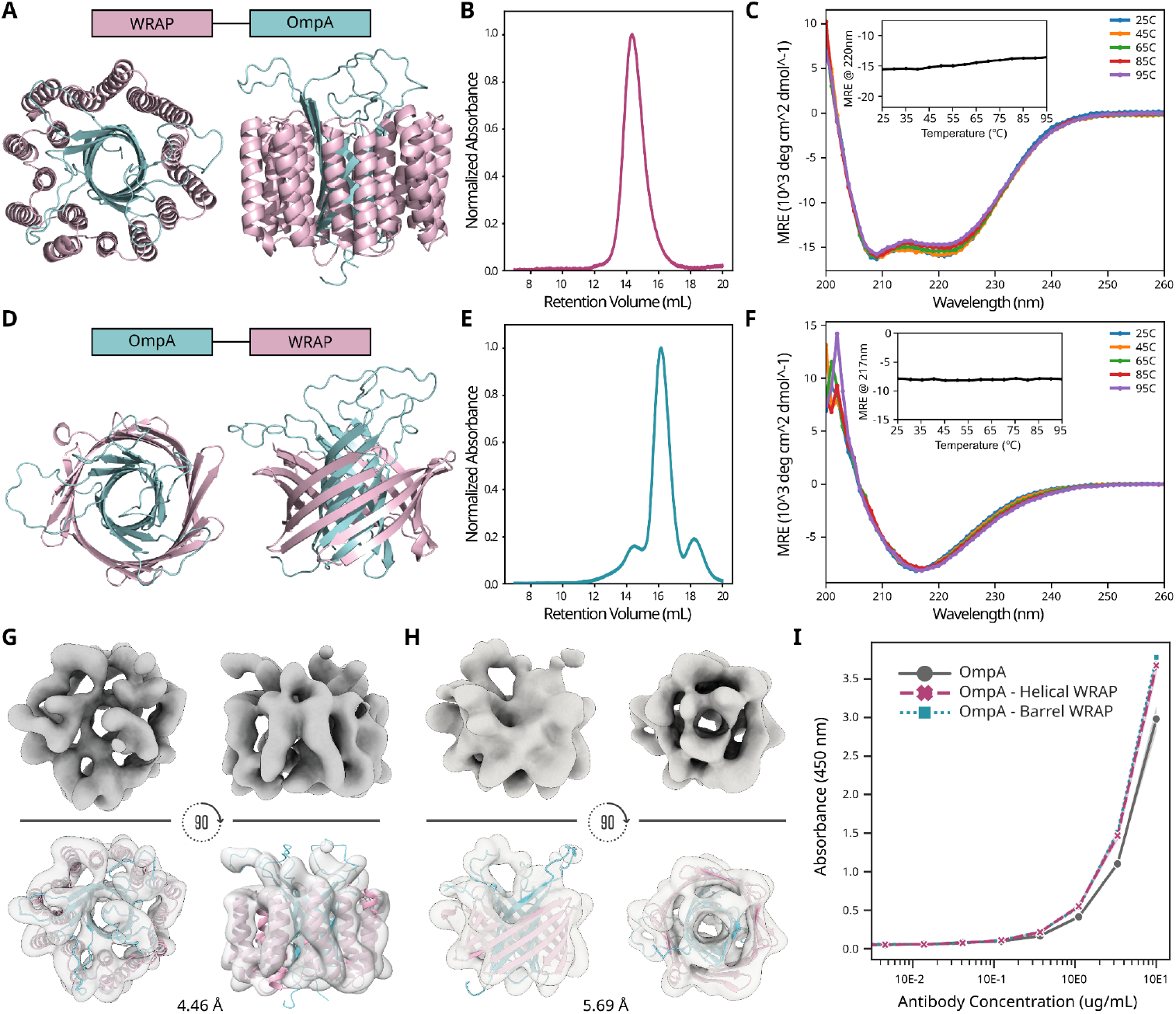
Solubilization of OmpA using helical and beta-barrel WRAPs. **(A, D)** Design model showcasing helical-and beta barrel-WRAPed (pink) OmpA (cyan) designs. **(B, E)** SEC traces of purified protein from soluble fraction. **(C, F)** Circular Dichroism (CD) data validating the thermal stability of WRAPed OmpA. **(G**,**H)** CryoEM data for helical and barrel WRAP-OmpA. CryoEM 3D density map and the computational design rigid-body docked into the density map showcasing high agreement to the design. **(I)** ELISA results confirming the binding of polyclonal OmpA antibody to the helical and barrel OmpA WRAP.

We used cryo-electron microscopy (CryoEM) to characterize the structures of both alpha-helical and beta-barrel WRAPed OmpA. CryoEM 3D reconstructions at 4.46 Å resolution for the helical WRAP-OmpA and 5.69 Å for the beta-barrel WRAP-OmpA were consistent with the intended locations of secondary structure elements of both the native OmpA beta-barrel and the surrounding designed WRAP (Figure 2G, 2H). Both WRAP structures highlight the generalization capabilities of RFdiffusion as there are very few concentrically nested beta barrels in the PDB (none that are asymmetric monomers) and there are no significant hits (TM-score > 0.6 (19)) that align to the WRAP-OmpA structures from searches against all available structure databases using Foldseek (20). To assess the preservation of antibody recognition elements in the WRAP constructs, we carried out ELISA with a polyclonal OmpA antibody and found that the WRAPs retained the corresponding epitopes, when compared to OmpA purified with detergent (Figure 2I).

These data establish that our computational pipeline can generate custom alpha-helical or beta-barrel WRAP domains that result in soluble, hyperstable versions of target transmembrane proteins. Since the beta-barrel WRAPs required significantly more computing to achieve similar AF2 metrics (Figure S1B), we focused primarily on designing helical WRAPs for the remainder of our targets.

## WRAPed helical multi-pass transmembrane proteins are functional

We next sought to solubilize helical membrane proteins while retaining their functionality. GlpG, a rhomboid protease located in the inner membrane of *E. coli*, is one of the best biochemically characterized intramembrane proteases and cleaves its substrates directly inside the lipid bilayer (21). Employing our WRAPs pipeline, we succeeded in solubilizing both the GlpG enzyme (Figure 3A) and its catalytic mutant (S201A+H254A) (22). Two out of the six WRAP-GlpG designs were detected in the soluble fraction of *E. coli* lysates and eluted at the expected retention volume during SEC. Representative SEC traces are shown in Figure 3B for the WT and in Figure S1 for the catalytic mutant. CD indicated that WRAPed GlpG, like OmpA, was highly thermostable, retaining its secondary structure even when heated to 95°C (Figure 3C). By contrast, native GlpG solubilized in detergent has a transition midpoint temperature of 77°C (22). CryoEM 2D class averages revealed clear secondary structural elements that align well with the computational design model (Figure 3D). Additionally, rigid-body docking of our computational design model into a cryoEM 3D reconstruction obtained at a resolution of 4.74 Å demonstrated strong agreement between the two (Figure 3D). To evaluate the activity of WRAP-GlpG, we used a fluorophosphonate serine hydrolase probe, FP-TAMRA, which selectively labels functionally active enzymes. Time-dependent active probe labeling showed that WRAPed GlpG accumulated substantial labeling over a two-hour period, while labeling of the catalytic mutant was nearly undetectable (Figure 3E). Together, these data show that WRAPs can not only preserve protein folds, but also help maintain functional integrity and enhance the thermostability of the solubilized membrane proteins. These results also show that the WRAP approach can be applied to the second major class of membrane proteins, helical multi-pass membrane proteins.

**Fig. 3.**
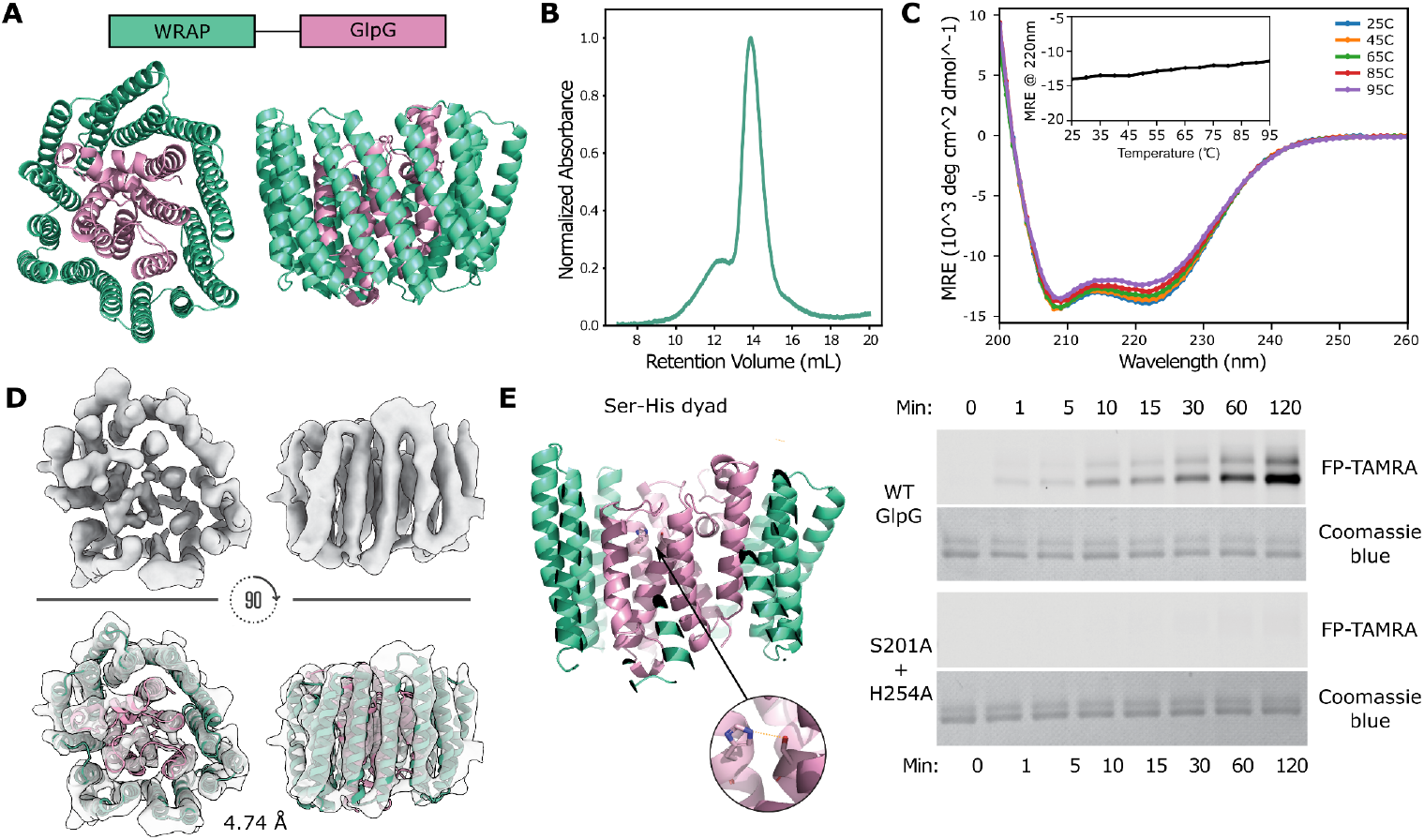
WRAPed intramembrane protease GlpG maintains activity. **(A)** WRAP (in green) and GlpG (pink) design model. **(B)** SEC trace of GlpG protein purified from the soluble fraction of *E. coli* lysates. **(C)** CD data shows that WRAP-GlpG is thermostable. **(D)** CryoEM data for WRAP-GlpG. CryoEM 3D density map and computational design model rigid-body docked into the density map, revealing high agreement. **(E)** From left to right: catalytic Ser-His dyad of WRAPed GlpG. Time-dependent FP-TAMRA active probe labeling of WT WRAP-GlpG in comparison to the catalytic mutant. Coomassie blue stain shows the total protein content in each reaction.

## Solubilizing *Treponema pallidum* outer membrane proteins

Targeting OMPs in vaccine development primes the immune system to combat pathogenic bacteria like *Treponema pallidum* (*Tp*), the causative agent of syphilis (23). To date, there are no reported experimentally determined structures of *Tp* OMPs (23), and the production of wild-type proteins from *E*.*coli* has been challenging. Because of their potential as vaccine antigens, we aimed to solubilize four putative *Tp* OMPs: TP0126, TP0479, TP0698, and TP0733 (3,23). These proteins are 8-stranded beta-barrels with transmembrane domains predicted with high confidence by AF2 (pLDDT > 90). Starting from these models, we employed a strategy similar to that described above for OmpA, and succeeded in solubilizing all four *Tp* OMPs. Following expression in *E. coli*, 19 out of 58 TP0126-wrap designs, 3 out of 10 TP0733 designs, 7 out of 29 TP0698 designs, and 4 out of 24 TP0479 designs were readily soluble and eluted at the expected size during SEC purification. Designs with the most prominent SEC peaks were selected for further characterization, and representative design models for TP0733, TP0126, and TP0698 WRAPs along with their respective SEC traces and CD data are shown in Figure 4. We also attempted to solubilize *Tp* proteins by swapping our WRAPs with the human ApoA-I protein (without using any additional tags). *Tp* proteins expressed as single fusions with ApoA-I failed to produce SEC peaks of the correct size (Figure S1E). 2D cryoEM class averages of WRAP-TP0126 and WRAP-TP0698 indicate secondary structure elements matching those in the computational design models and AF2-predicted structures (Figure 4J, 4K). Low-resolution cryoEM 3D reconstructions (7.17 Å for TP0126 and 4.01 Å for TP0698) were consistent with the overall size and geometry of the WRAP-TP0126 design model and were in good agreement with the predicted structure of WRAP-TP0698 (Figure 4J, 4K and Figure S2D, S2E). The density corresponding to the extracellular loops of TP0698 differed from the AF2 prediction, which is not surprising as the prediction has very low confidence in this region (pLDDT < 50).

**Fig. 4.**
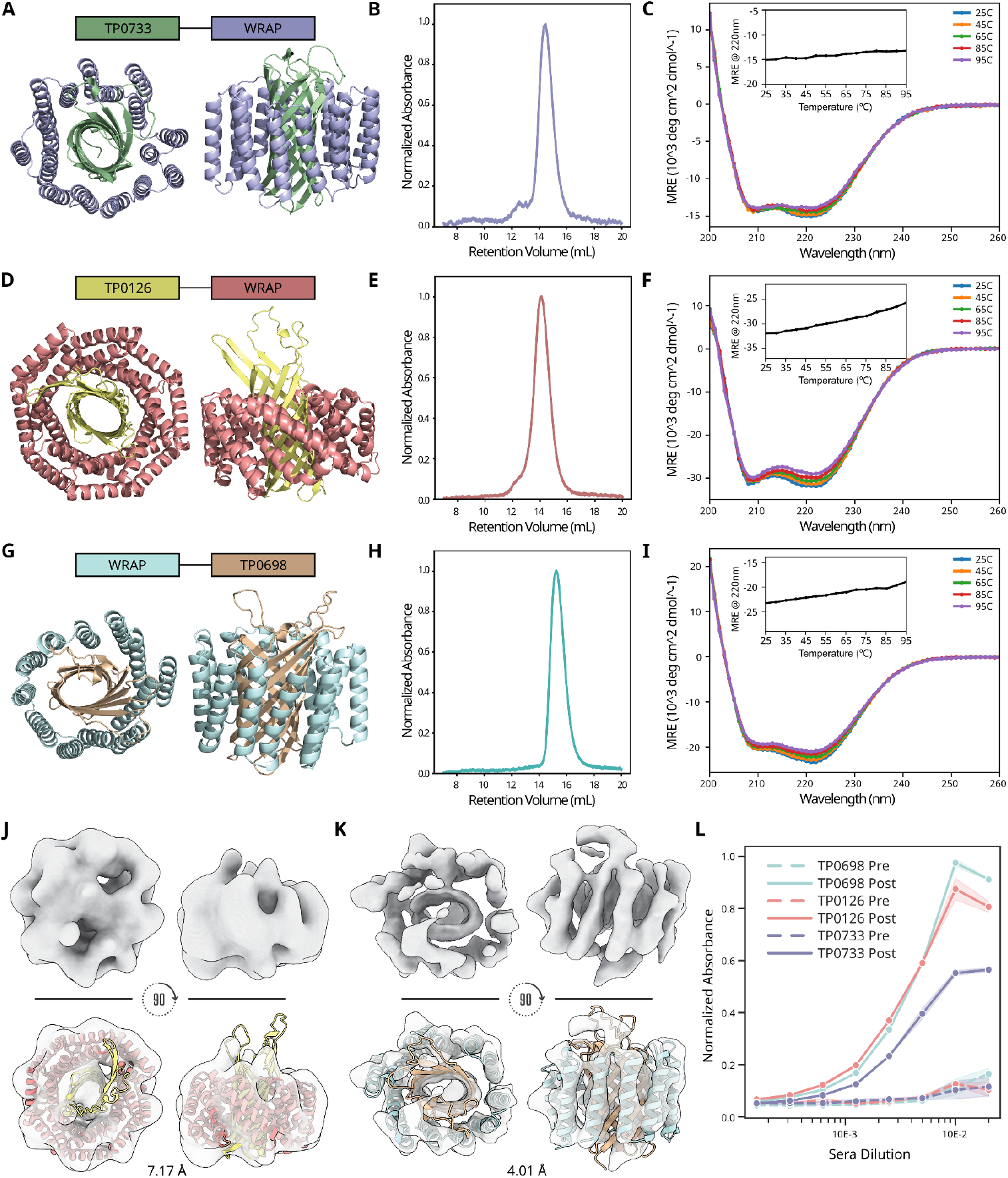
Solubilization of *T. pallidum* OMPs. **(A**,**D**,**G)** Design models of the TP0733, TP0126 and TP0698-WRAP. **(B**,**E**,**H)** SEC traces of purified TP0733, TP0126 and TP0698-WRAPs. **(C**,**F**,**I)** CD spectra of TP0733, TP0126, and TP0698-WRAPs demonstrating thermostability. **(J**,**K)** CryoEM data for TP0126-WRAP and WRAP-TP0698. 3D cryoEM density maps and computational designs rigid-body docked into 3D map highlighting agreement in the overall map size and shape compared to the design model. The extracellular loop domains of TP0126 are not resolved and the extracellular loop conformations of TP0698 appear distinct from the AlphaFold prediction. **(L)** ELISA results show specific response against WRAPed Tp OMPs with pooled serum from Kymouse Platform mice vaccinated with freeze-thaw inactivated *Treponema pallidum* subsp. *pallidum* strain Nichols.

To determine antigenic properties of WRAP-TPs, we used sera from humanized Kymouse platform mice and rabbits immunized with *T. pallidum*. Mice were immunized with freeze-thaw inactivated *Treponema pallidum* (24), and rabbits were challenged with live *Tp* spirochetes (25). ELISA experiments showed that the rabbit serum antibodies bound WRAP-TP0698 (Figure S2F), and the mouse serum bound all three WRAPed antigens, whereas there was no reactivity from pre-immune sera (Figure 4L). There are currently no available monoclonal antibodies targeting these antigens, as recombinant expression of the wild-type proteins in *E. coli* failed and thus there has been no way to directly immunize to generate them or to generate probes necessary to isolate them (scaffolds displaying single ECLs on a soluble scaffold do react with immunized rabbit serum (25), but such scaffolds cannot be used to detect antibodies that bind epitopes involving multiple extracellular loops or the beta barrel core). Current efforts are directed at identifying the antibodies that react with the WRAPed antigens. In this way, WRAP-TP antigens could help generate, isolate, and characterize monoclonal antibodies, and have considerable promise as components of a potential syphilis vaccine.

## Conclusions

We introduce a method for solubilizing native transmembrane proteins using genetically encoded designed auxiliary domains that requires neither detergents or refolding, enabling expression in the *E. coli* cytoplasm with enhanced stability while preserving native sequence, structure, and function. Unlike previous approaches (5–8), WRAPs are tailored to the membrane protein of interest, requiring no modification of the native sequence. Additionally, the sequences of our WRAPs are synthetic and do not have substantial sequence homology to known human proteins; this could be useful in the context of vaccination, where eliciting antibodies that target self-proteins (e.g., ApoA-I (9)) is undesirable. CryoEM 2D class averages and 3D map data demonstrate that soluble-WRAPed OmpA and GlpG proteins retain their structural integrity, as confirmed by comparing the resulting 3D maps with design models and previously deposited structures in the PDB. WRAP design does not require experimentally determined structures, as illustrated by the solubilization of syphilis outer membrane beta barrels starting from AF2 models. In addition to solubilizing integral membrane proteins for use as vaccine antigens, WRAPs could be useful quite generally for developing probes for traditionally challenging membrane protein targets, for example by facilitating the production of soluble, stable, and antigenically intact versions for monoclonal antibody generation or designed binder characterization. As illustrated by the production of soluble versions of extremely difficult to characterize syphilis outer membrane protein antigens, WRAPs should contribute to diagnosing and combating bacterial diseases which have been limited by the difficulty in producing and characterizing the major surface antigens.

## Acknowledgments

The authors would like to thank Matthew DeLisa for advice and encouragement. Special thanks to Edin Muratspahic, Susana Vasquez Torres, Yakov Kipnis, Anna Lauko, Sam Pellock and Arvind Pillai for helpful discussions. We would also like to thank Justin D. Radolf (UConn Health) for insight into and helpful discussions on *T. pallidum* outer membrane proteins and Stephen Reece and Jose Luis Slon Campos (Kymab, a Sanofi Company, Babraham Research Campus, Cambridge, UK).

## Author contributions

D.B, N.P.K., M.J.C, L.M., and D.E.K. conceptualized the project. L.M. and D.E.K. developed the design pipeline, with input from S.M and H.E.E. Authors L.M, D.E.K, P.B., and H.E.E. designed, screened, and experimentally characterized WRAPs. A.J.B., A.C., and C.W. obtained and processed cryo-EM data. M.L., A.M., R.R., E.C.W., and S.H. performed protein purification. Q.L. performed ELISA with mice sera. L.G. and N.T. provided the bacteria for the immunization work. X.L. performed LC/MS experiments. D.B., N.P.K., K.L.H, M.J.C., P.C. and L.S. provided supervision and funding. L.M., D.E.K., H.E.E., N.P.K., and D.B. wrote the manuscript. All authors discussed the results and commented on the manuscript.

